# Structural basis of *Drosophila* insulin receptor activation by DILP2 hormone

**DOI:** 10.1101/2025.08.22.671756

**Authors:** Talha Shafi, Olga V. Moroz, Huw T. Jenkins, Maria Chechik, Michail N. Iusupov, Marta Lubos, Jiří Jiráček, Pierre De Meyts, Andrzej M. Brzozowski

## Abstract

Insulin-related hormones regulate key life processes in the animal kingdom, from metabolism to growth, lifespan and aging, through an evolutionarily conserved insulin and insulin-like hormones signalling axis (IIS). In humans the IIS axis is controlled by insulin, two Insulin-like Growth Factors, two isoforms of the insulin receptor (hIR-A and -B), and its homologous IGF-1R. In *Drosophila*, this signalling engages seven insulin-like hormones (DILP1-7) and a single receptor (dmIR) that follows the blueprint of hIR/hIGF-1R. This report describes two cryo-EM structures of the dmIR ectodomain dmIR-ECD:DILP2 complex, revealing their structural homology with dmIR:DILP5 complex. The high excess of DILP2 yielded two dmIR-ECD complexes in asymmetric conformations, similar to that observed in some complexes of hIR and in the dmIR-ECD:DILP5 complex. This stoichiometric and structural heterogeneity, yielding one- and two-DILP2:receptor complexes – were not observed in DILP5:dmIR-ECD assembly. Also, the resistance of dmIR-ECD to form more DILP2 saturated complexes, despite very high excess of this hormone, suggest that the specificities of DILPs may lie in their *k*_on_/*k*_off_ kinetic parameters. This work expands understanding of the dmIR conformational flexibility, suggesting also that insect dmIR follows more hIR rather than hIGF-1R receptor signal transduction pattern induced by various DILPs.

## INTRODUCTION

The insulin/insulin-like growth factor signalling axis (IIS) is an evolutionarily ancient, highly conserved, endocrine and paracrine signal transduction network in the animal kingdom: from freshwater polyps to humans (1, 2). Its pleiotropic spectrum spans such key life processes such as metabolism, growth, development, reproduction, aging and lifespan, and cell growth, differentiation and migration.

In humans, IIS is govern by insulin secreted by pancreas β-cells, and Insulin-like Growth Factors-1/2 (hIGF-1/2) produced in multiple tissues. Insulin and hIGF-1/2 signal through closely related cell-surface receptor tyrosine kinases (RTKs), the insulin and IGF-1 receptors (hIR, hIGF-1R), and through their hybrid dimers (3-5).

Insulin acts through two closely related (12 amino acid difference) hIR isoforms: hIR-A - also with some mitogenic properties, and hIR-B that shows mostly metabolic effects, while hIR-A:hIGF-1R hybrids increase the complexity of IIS (6). Insulin homologous hIGF-1/2 are not stored in crystalline/aggregated forms as insulin, but are secreted by multiple tissues as monomers, maintained in biological fluids in complexes with several IGF binding proteins (IGFBP1-6) that regulate their bioavailability (7).

HIR and hIGF-1R are a unique Class II of RTKs family being -SS-bridges linked (αβ)_2_ subunits homodimers – even in their ligand-free, apo forms, with the same modular, exons-reflecting organisation (**Fig. 1a**)(8). In hIR (and hIGF-1R) the α-subunits consist of L1 (Leucine-rich repeats domain 1), CR (Cysteine Rich), L2, FnIII-1 (Fibronectin type III-like), FnIII-2 and alternatively spliced ID (Insert domain) modules (5, 9) The β-subunit starts with the remaining part of the spliced ID/FnIII-2 domain, followed by the FnIII-3 domain, the transmembrane helix (TM), juxta-membrane (JM) and TK modules. Both α-subunits are linked by several structurally and functionally important -SS-bonds, while a single -SS-bond connects α–β-subunits. Human IR (and m(mice)IR) structures consistently reveal the spectrum of conformations from the apo-IR in an inverted V - Λ-like - form, to a very different - fully four-insulin saturated, almost symmetrical T-shaped form with insulins in four sites. However, many ‘intermediate’ hIR/mIR structures have been found between these two opposite IR conformers, with one, two, or three insulins bound (**Fig. 1b**) (4, 5, 10-16). These intermediate structures have the α-subunit L1-CR-L2 arms of the receptor in many asymmetrical *Γ* and *T* conformations, from one arm-down/one-up to the different stages of detachment of the down-arm from the FnIII-1–FnIII-3-stem of the receptor.

**Figure 1.**
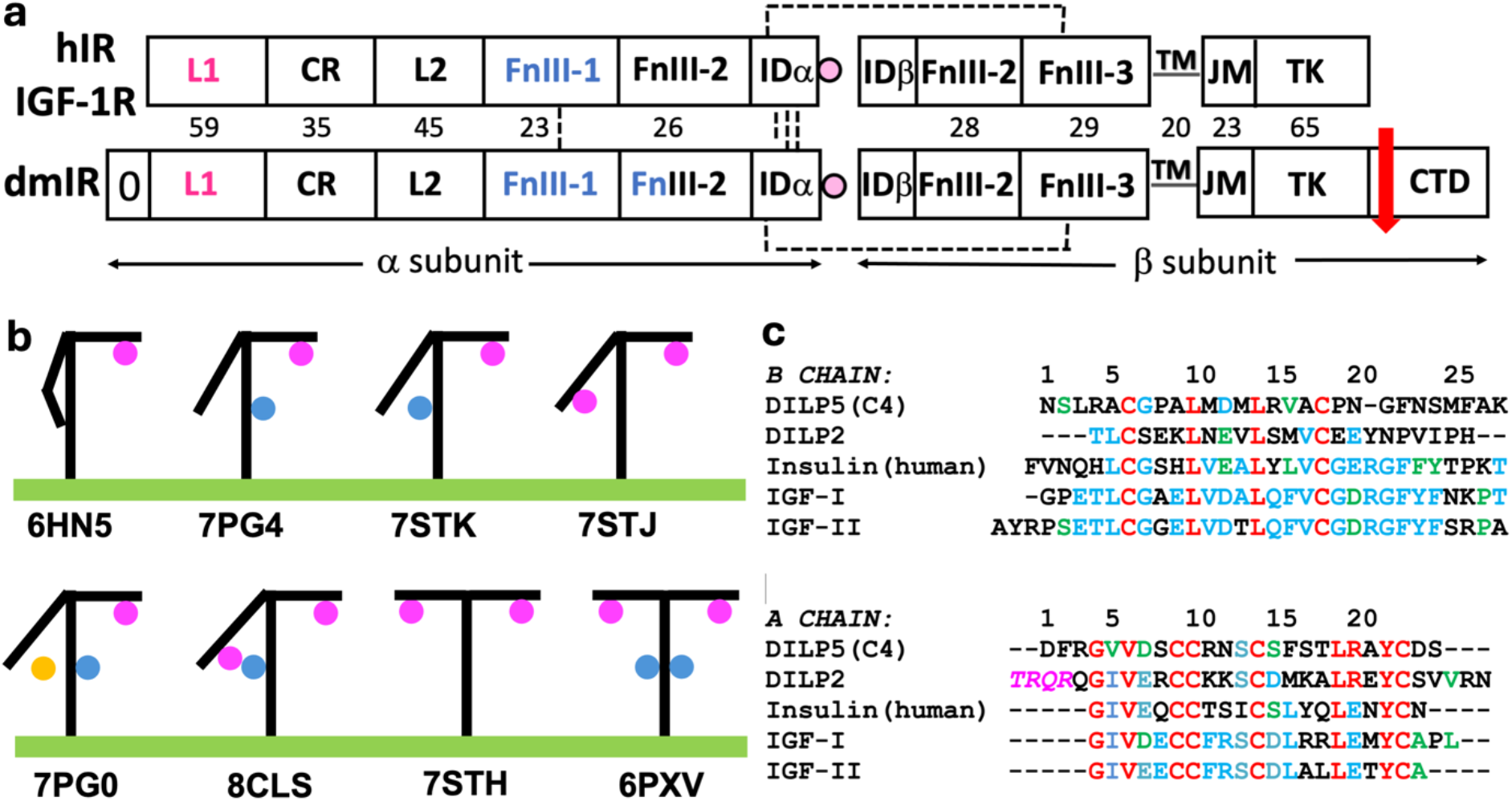
**(a)** Domain organisation of the hIR/hIGF-1R (top) and dmIR (bottom)(not up to scale). Domains contributing to hormone binding site 1 are in magenta, and to site 2 in blue. The dash lines indicate disulphide bonds (one less between dmIR ID domains than in hIR/hIGF-1R). Red arrow stands for *in vivo* processing of the dmIR C-Terminal Domain. **(b)** Schematic depiction of the representative conformers of Insulin-like Receptors ECDs; hormone (circle) bound to site 1 in magenta, to site 2 – in blue, in-between site 1**’**-2 - in yellow; cell membrane – green line; PDB IDs for some typical conformers are given below the membrane. **(c)** Sequence alignment of DILP5 (C4 variant), DILP2, human insulin, IGF-I and IGF-II: IGF-1/IGF-II additional B-A chain linking C-domains and C-terminal D-domains are omitted for clarity. DILP2 A chain N-terminal ThrA1-ArgA4 (in magenta and italic) were not included in synthesis of this hormone.

In contrast to a limited number of insulin-like proteins (ILPs) in humans (insulin, hIGF-1/2), such hormones are very diverse in the animal kingdom, ranging from 7 two-chains ILPs in *Drosophila* (DILP1-7), to 40 ILPs in *Bombyx mori* and *Caenorhabditis elegans* (17-19); (*Drosophila* DILP8 is a paralogue of human relaxin binding to a G-protein-coupled receptor, Lgr3 (20)). Nevertheless, all these hormones share similar two A, B-chains motifs (with an extra C-domain in some single-chain ILPs), linked by inter-/intra-chain disulfides. Despite the variety of the ILPs in invertebrates they act in *Drosophila melanogaster* (Dm) and *Caenorhabditis elegans* through a single so-called insulin receptors (17, 19, 21). This multiplies the functions of *Drosophila* insulin receptor (dmIR) into many life processes including development, regulation of life span, metabolism, stress resistance, reproduction and growth, to assure an efficient control of insect’s life cycle (22-24).

We have recently determined the first cryo-EM structure of an invertebrate IR: the ectodomain of *Drosophila melanogaster* dmIR (∼33% sequence identity in the hIR overlapping regions (25, 26)), in the complex with DILP5 hormone (dmIR-ECD:DILP5) (27). This tilted dmIR *Γ* -like conformer, with three bound DILP5 molecules, revealed a remarkable similarity of dmIR and hIR/hIGF-1R 3-D organisations on all structural - quaternary to secondary - levels. The strikingly similar structural outlines of these receptors were also reflected in their similar hormone:IR binding modes.

The two-chain DILP5 in the dmIR-ECD:DILP5 complex is bound to the main hormone binding site 1 on the *Γ* – higher arm of the receptor – in the same fashion as hormones in h(m)IR/hIGF-1R systems. In all these IR-like receptors the main, so-called high affinity site 1 is being formed by the L1 domain and the crossing-over α–CT**’** terminal region of the ID**’** domain from the other α-subunit (the (**’**) denotes contribution from the other symmetrical αβ half of the IR, i.e., there are potentially two sites 1: L1/α–CT**’**, L1**’**/α–CT)). The α–CT**’** segment lie across the L1β2 sheets of the L1 domains, shifting from its apo-IR position upon ligand binding to mediate a mostly indirect tethering of insulin/hIGF-1/2/DILP5 onto their respective L1β2 surfaces. The so-called insulin low-affinity sites 2/2**’** are on surfaces of the FnIII-1/FnIII-1**’** domains has been occupied by hormones in many hIR complexes, except hIGF-1R where only one site 1 is engaged by hIGF-1 (or hIGF-2) (11, 28, 29). Interestingly, site 2 in the dmIR-ECD complex, that is close to the lower arm of the receptor, is occupied by DILP5 in a slightly different manner than in the hIR complexes. Moreover, this hormone molecule is in close contact with the third DILP5 which sits in site 1’, giving these two (site 2/1’) bound DILP5s a hitherto unseen arrangements of the ligands.

The structural conservation of TK Insulin-like Receptors in *Metazoa* is also highlighted by their inter-species functional homology. For example, dmIR can be activated by human insulin (30, 31), DILP5 shows the insulin-like bioactivity in mice (31), and cone snail venom insulin activates hIR (15).

Studying *Drosophila* IIS contributes to the understanding of the fundamentals of this key signaling network, but it also has an important translational aspect, as this fruit fly is a widely used model for many human pathologies - from metabolic disorders to neurodegeneration - with over 75% of human-disease-related genes are conserved in this animal (32, 33).

Despite astonishing 3-D insights into the holo- and apo forms of hIR/hIGF-1R, mIR, and, recently, dmIR, several key questions about their insulin-binding triggered activation, and complex allosteric signal transduction, still remain. The recently reported unexpected proximity of dmIR-ECD:DILP5 site 1**’**/site 2 bound hormones expanded the broad spectrum of IR-like receptor stoichiometries and conformation (5, 27). It also prompted a question of mode of binding of other DILPs: what are their unique signatures and common conformational features.

Therefore, we focused in this work on deciphering the nature of dmIR interaction with another DILP hormone: DILP2 (**Fig. 1c**). Despite some compensatory functional overlapping with other DILPs (34), DILP2 seems to be one of the most prevalent and physiologically important DILPs. It is expressed in Insulin Producing Cells (IPC) in brain (as DILP5), but also in Imaginary Disks, salivary glands and Glial cells of the CNS, being important for trehalose metabolism, stress response, life-time fecundity and other physiological effects (35). Interestingly, on a signaling level, DILP5-dmIR interactions results in a high, continuous phosphorylation of the key downstream effector Akt, while such response is very transient for DILP2 (36).

Here were report two cryo-EM structures of dmIR-ECD:DILP2 complex. Interestingly, in contrast to the dominating three-DILP5:one-dmIR conformer (27), we observe here a heterogeneity of DILP2 complexes that allowed description of one- and two-DILP2-bound - dmIR-ECD assemblies.

## RESULTS

The 291Leu-1309Ala dmIR-ECD construct with C-terminal StrepII-tag has been expressed in the *Sf9* cells baculovirus system, and purified on the Strep Trap HP affinity media, and by the size-exclusion chromatography as described before (27). In contrast to DILP5, the determination of the DILP2:dmIR-ECD *K*_d_ by the μITC was difficult, with titration curves showing none-or-all binding, indicating likely a very low DILP2 affinity to dmIR-ECD. DILP2 poor receptor binding properties were also reflected in a very high 40:1 hormone:dmIR-ECD ratio that was necessary to obtain workable cryo-EM grids; lower hormone excesses gave grids with an overwhelming presence of disordered/apo forms of the ectodomain. It should also be noted that for the feasibility of the full synthesis the DILP2 A-chain starts here at Gln(Q)A5; the TRQR N-terminal A1-4 residues were omitted in the synthesis, also due to still uncertain hormone’s *in vivo* post-translational processing of these regions (as for DILP5 (31)). The full physiological relevance of such DILP2 construct had already been confirmed (37), but the numbering of the DILP2 A-chain in this paper follows ThrA1 as this chain N-terminal residue, to align with the UniProtKB DILP2 entry (ID: Q9VT51).

The alternative analysis of the subset of particles allowed determination of cryo-EM structures of the two main tilted *Γ* forms of the dmIR-ECD:DILP2 complexes. The main form 1, referred here as to conformer **1**, contains only one DILP2 hormone in the classical, upper arm, site 1. The minor form 2, referred here as conformer **2**, has similar DILP2-occupied upper arm site 1, but the maps indicate also a likely presence of another DILP2 here, wedged between the lower arm site 1’ and the stem of the ectodomain around FnIII-1 site 2 region.

### The structure of conformer 1

The overall structure of the form 1 of the DILP2 complex is, overall, very similar to the dmIR-ECD:DILP5 assembly, showing tilted *Γ*-like conformation (**Fig. 2a**). The upper arm of the complex contain classical site 1-bound DILP2, while the ligand-free lower arm (e.g., L1**’** domain) moves closer to the FnIII-1/FnIII-2 stem of the receptor by ∼9 Å (if the upper arms site 1-containing dmIR αβ subunits of DILP5 complex and DILP2 conformer **1** are superposed on the Cα atoms). This movement is more significant for the CR**’**, and, especially, L1**’** domain, while L2**’**/FnIII-1**’**/FnIII-2**’** folds follow rather closely their DILP5 complex orientation. Interestingly, despite only one bound-hormone, the lower arm of conformer **1** is not touching the stem of the receptor leaving around 20 Å gap there, making it, possibly, a more unique signature of conformer **1** among insulin-like hormones:receptor complexes (**Fig. 2a**). An unoccupied gap between the lower arm and the stem of the IR is usually associated with the presence of more insulins, e.g., like in the PDB ID 7PG4 complex (even at hormone unsaturated conditions) (16), where such - much bigger than in conformer **1** - gap exists at the lower arm of the hIR; however, the other half of this receptor - site 1/2**’ -** are occupied there by insulins. One insulin:hIR - or hIGF-1:hIGF-1R - complexes show a good occupancy of higher arm site 1 (e.g., (10, 38), while the receptors’ lower arms in these complexes are in close, touching, contact with their fibronectins stem. Therefore, such low occupancy of the dmIR-ECD *Γ*-like conformer, and its single hormone:receptor-like conformation obtained at 40:1 hormone’s excess are rather unexpected, and may represent some physiological relevance (see later).

**Figure 2.**
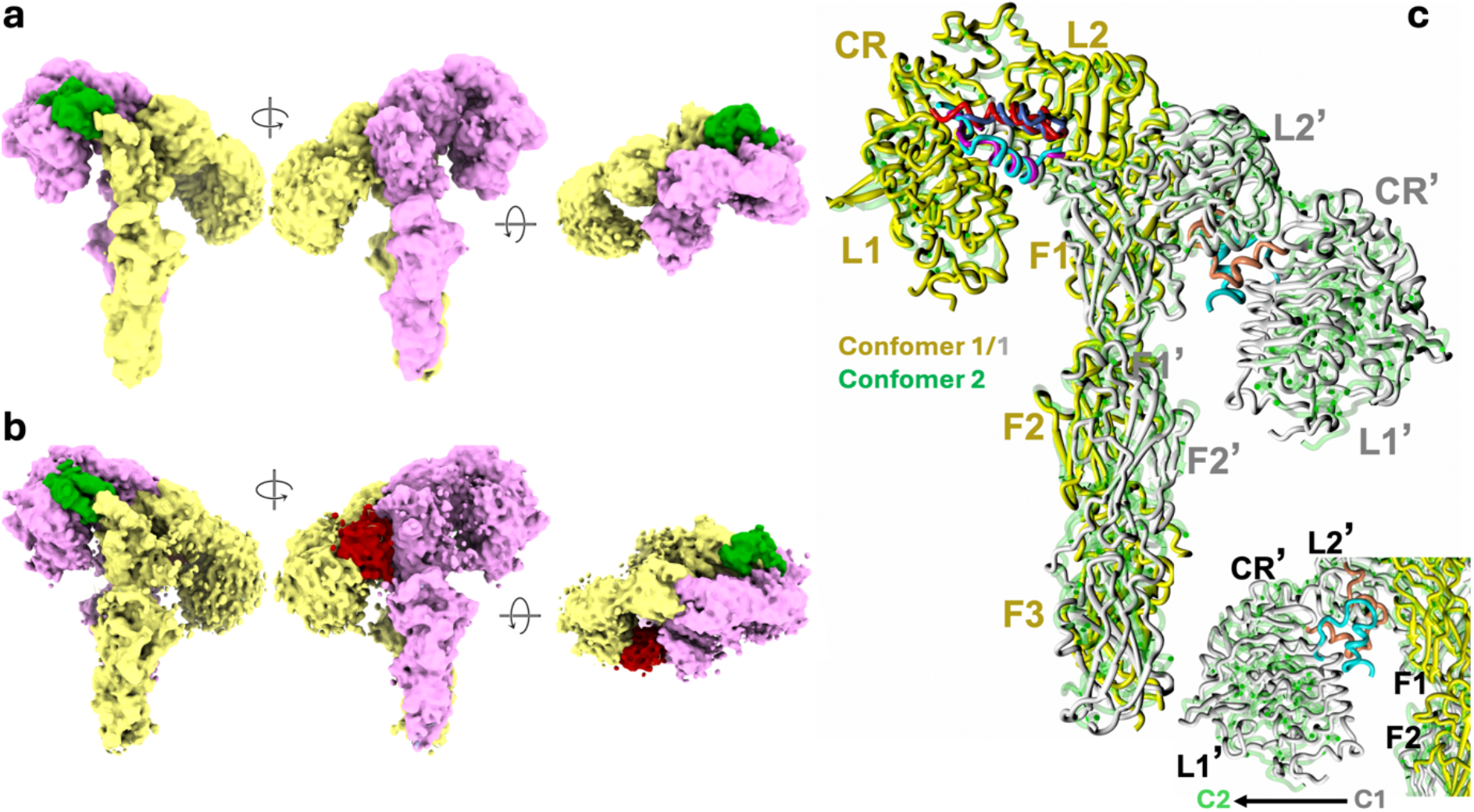
Surface representation of cryo-EM maps for DILP2 conformer **1 (a)** and **1 (b).** The lower arms are yellow and the higher ones in magent. DILP2 in sites 1 are in green, and in dark red in between sites 1**’**-2 in conformer **2. (c)** SSM superposition (see Methods) of the upper arms receptors’ conformer **1** (in yellow) and **2** (in green); the lower arm of conformer **1** is in grey – conformer **2** also in green. Individual domains are indicated by single letter abbreviations, i.e. FnIII-1 = F1, etc. DILP2 site 1 B chain of conformer **1** is in blue – conformer **2**: in purple, A – chains there are in red and deep blue, respectively. The B-chain of the in-between sites 1**’**-2 DILP2 in conformer **2** is in blue, its A chain in coral. The bottom insert shows the other view (180^°^ rotated from the upper image) of the ‘bottom’ DILP2 in conformer; the shift of the lower arm from conformer **1** to **2** is indicated.

The overall arrangement of DILP2:site 1 in conformer **1** is similar to DILP5 binding mode, reflecting conserved tethering of insulins onto L1β2 surface *via* α-CT**’** helix (**Fig. 3a**). DILP2 three amino acids shorter B-chain N-terminus fits well between body of this hormone and FnIII-1 loops, with the ThrB1 making likely hydrogen bonds to Glu833. As the DILP5 B1-B3 residues that filled the B-chain N-terminus channel are missing in DILP2, the Thr827-Ile842 loop (especially its Pro833-Pro837 section) - postulated gatekeeper of this region - moves closer over this space, indicating conformational dynamics of this loop that may indeed depend on the nature and length of DILPs’ B-chain N-termini. The ProB8→GluB5 DILP5/DILP2 change does not affect the beginning of the B5-B11 α-helix. It is still tethered onto α-CT**’** segment and L1β2 surface by mostly hydrophobic interactions observed in the DILP5 complex, while DILP2 contacts are supplemented there by AsnB8-Tyr392 hydrogen bond, resulting from MetB11→AsnB8 mutation in DILP2 hormone. It seems that a more substantial structural change between DILP2 and DILP5 in site 1 occur at the SerB12/ArgB15-AsnB19/PheB22 turn. This turn runs closer to the core of DILP2 hormone, and the hydrophobic cavity that is quite isomorphous in human insulin and DILP5 complex being filled by PheB24 and PheB22, respectively, it is occupied by AsnB19 in DILP2 (**Fig. 2a**). Also, DILP2 TyrB18, that could – potentially – fill this cavity, is pointing into solvent. The other important receptor hydrophobic cavity:hormone B-chain interaction - maintained in human insulin by PheB26 and MetB25 in DILP5 complexes - is fulfilled in DILP2 by IleB22. In general, the classical (in insulin:hIR system) B22-B26:receptor interface that is well mimicked in DILP5:dmIR-ECD complex, it seems to be less favourable/weaker in the DILP2:dmIR-ECD assembly. The additional – to DILP5 and human insulin – A-chain N-terminal (ThrA1-GlnA3) and C-terminal (ValA27-AsnA30 are not visible in the maps hence they are not likely strong contributors to DILP2 site 1 in conformer **1**.

**Figure 3.**
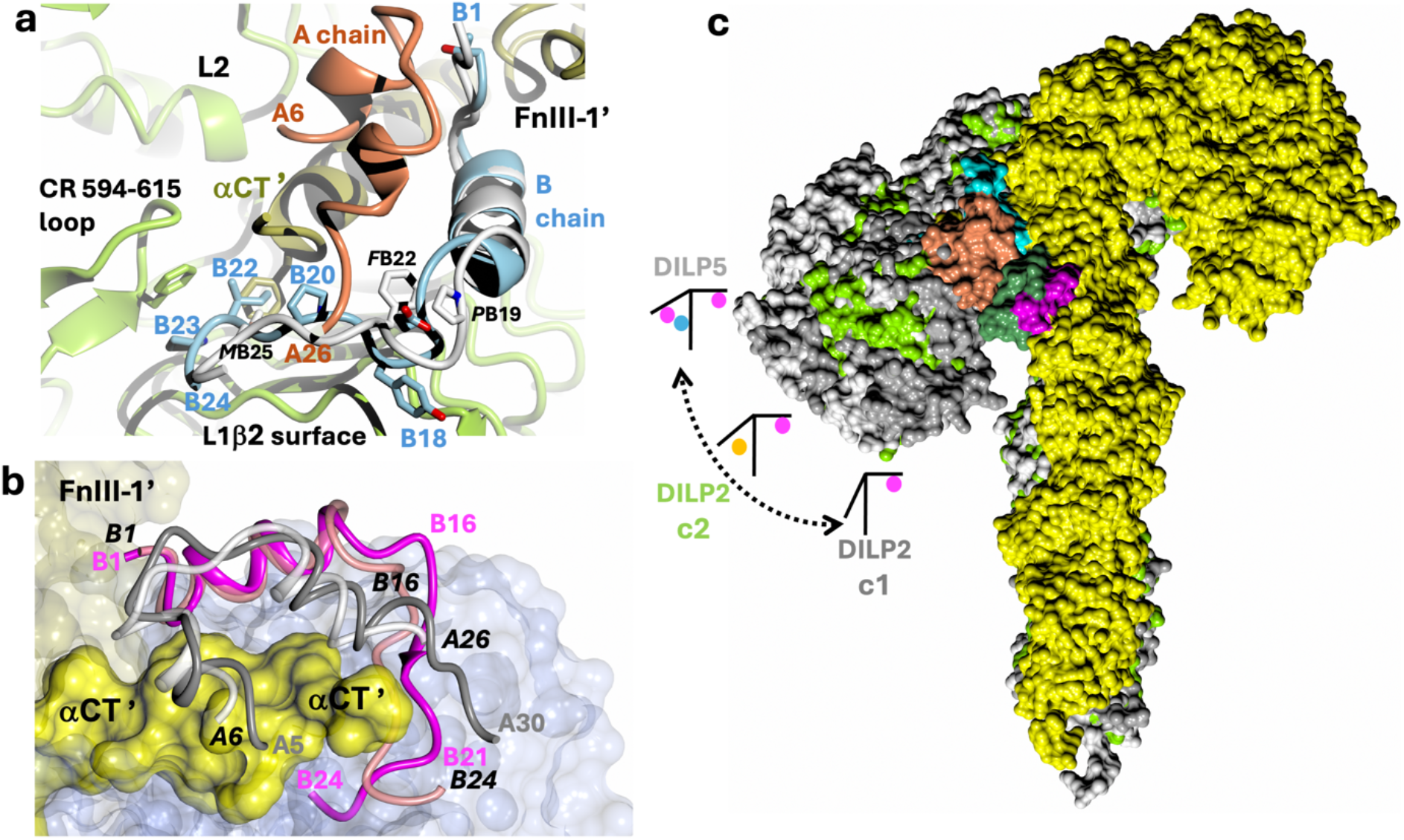
**(a)** An overlap of dmIR-ECD upper arm site 1 in DILP2 and DILP5 complexes. The main elements of the dmIR-ECD site one are in green. DILP2 B-chain is in blue, and its A-chain in coral, while DILP5 B-chain is in white; DILP5 A-chain is omitted for image clarity. DILPs were superposed on the B5-B14 Cα atoms. Labels for some DILP2 B-chain residues are in blue, and in black for DILP5. Only side chains of some key residues discussed in the main text are shown. **(b)** On overlap of B-chains (on the B5-B14 Cα atoms) of DILP2s in site 1 in conformer **1** (coral) and conformer **2** (magenta); DILP2 conformer **1** A-chain in white/grey, A-chain of conformer **2** – dark grey. The site 1 of conformer **1** is shown as surfaces, with α-CT**’** as yellow spheres. **(c)** Surface representation of upper-arms superposition of DILP2:dmIR-ECD conformer **1**, conformer **2** and DILP5:dmIR-ECD complexes. The upper arm αβ subunit with of DILP2 conformer **1** is in yellow, its lower arm αβ subunit in dark grey; both (αβ)_2_ dmIR-ECD subunits of DILP2 conformer 2 are in green, and those of DILP5:dmIR-ECD complex are in white. B chain of site 2 DILP5 is in magenta, and its chain A is in dark green; in coral – B-chain of DILP2 that is between sites 1**’** and 2, it’s A-chain in cyan. The lower arm site 1**’**-bound DILP5 from the DILP5:dmIR-ECD complex is invisible here being occluded by DILP2 from conformer **2**. The ball and stick diagrams (colour coding as in **Fig.1b**) show the transition-migration of the lower dmIR-ECD arm from no-DILP (DILP2 conformer **1** complex), one in-between DILP (DILP2 conformer **2** complex), and two DILPs (DILP5 complex) hormone:receptor assemblies.

### The structure of conformer 2

The overall structure of conformer **2** follows – in general – a tilted *Γ* form of the receptor, with its upper arm holding DILP2 in site 1. However, there is an extra density between the lower arm and stem of the receptor that indicates there the presence of a second DILP2 molecule (**Fig. 2b**).

DILP2 B-chain N-terminus in upper site 1 is accommodated similarly to its conformer **1** fold. However, ThrB1 seems to make different pattern of hydrogen bonds, and, interestingly, the longer in dmIR than in hIR gatekeeper Thr827-Ile842 loop moves away from the B-chain N-terminus channel – opposite to what is observed in conformer **1**. This suggest a more dynamic role of this loop in discrimination of different lengths DILPs B-chain N-termini, rather than its static – e.g., conformation stablising – functioning. The main difference between site 1 DILP2 binding mode in conformer **1** and **2** seems to occur in B15-B25 regions of the hormone. The fold of the B15-B18 turn is different in both conformers, and leads to +1 shift of register in conformer **2**. This results in lack of occupancy of ‘PheB24 hIR’ pocket and closer run of the B18-B21 chain to the core of the hormone (**Fig. 3b**). AsnB19 that fills in ‘PheB24’ pocket moves forward along the B-chain, being involved in likely hydrogen bond with NH-of Val1045 of the α-CT**’** segment. The more ‘tighter’ (to the L1 domain) run of the B18-B21 chain can result from more visible A-chain C-terminal A27-A30 residues in conformer **2**, which seem to be pushing the B-chain towards the receptor, running outside of the DILP2 molecule but without any particular strong interactions. However, the most remarkable difference in this region concerns the direction of the DILP2 B-chain C-terminal B21-B24 peptide, for which the maps are rather ambiguous but allow some alternative modelling of this Instead of pointing outwards of the L1 domain – as (what can be envisaged) in conformer **1** – it bends after ProB20 at ValB21 inside: over the L1β2 surface, fitting between this β-sheet and α-CT**’** segment (**Fig. 3b**). It can be accommodated there quite easily, without any steric clashes, with maybe a putative HisB24Nε2-Glu656 hydrogen bond, and series of hydrophobic-like interactions. It is interesting that, despite such unexpected turn at ValB21, the IleB22 occupies the same ‘PheB26’ cavity as in conformer **1**. It remains unclear whether such unusual – still putative, considering low quality of maps there - conformation of the B20-B24 is specific for the DILP2 hormone (e.g., being facilitated by two Proline residues there (ProB20, ProB23)), or it reflects its more dynamic character. It must be stressed here, however, that conformer **2** has been determined with an overall much lower resolution than conformer **1**, and although it has been fitted there with due care, its more unexpected structural features in site 1 should be considered with caution. Nevertheless, the alternative, quite different folds of the B20-B24 chains in DILP2 site 1 may not be surprising in the context of a very low DILP2 affinity for this dmIR-ECD construct. Such weak binding can result from (i) loss of filling of ‘PheB24’ cavity, (ii) weaker filling of the ‘PheB26’ cavity by IleB22 (instead of MetB25 in DILP5), (iii) dynamic character of B20-B24 chain that can interfere with hormone effective settling in the dmIR site 1, and from (iv) possible – alternative - steric hindrance of the B20-B24 L1β2 ‘inward’: intercalated between L1β2:α-CT**’** conformation, hence, subsequently, its weakening effect on this crucial interface.

The CR Ala594-Asn615 loop, which was suggested as a discriminatory feature that prevents binding of a single-chain like hormones (e.g. IGF-1/2), remains in the same conformation in site 1 in both conformers **1/2** and DILP5 complex, confirming its important structural and functional role (**Fig. 3b**).

The other main structural feature of conformer **2** is an additional density between lower arm L1**’** domain and FnIII-1 domain (**Fig. 2b**). Although a low resolution and poor quality of maps there do not allow an unambiguous positioning of DILP2, the occurrence of the hormone molecule in-between sites 1**’**-2 is rather obvious. The orientation of the lower arm in conformer **2**, and the presence of DILP2 there, is, overall, similar to the one observed in the PDB ID 7PG0 structure of hIR, which, however, has also third hormone bound to site 2**’** (16). Hence hormone:receptor arrangement seen in conformer **2** seems to be novel, expanding conformational landscape of the IR-like receptors. The orientations of hormones at this lower arm site in conformer **2** and in the 7PG0 are different, but they cannot really be compared as DILP2 maps are not unequivocally defined. The orientations of the lower IR arms – especially CR-L1 domains – are slightly different in conformer **2** and 7PG0 structure, with the latter one showing more hormone-embracing conformation. It is worth mentioning that in both of these structures the α-CT segments are not observed in the L1**’** domain on the L1β2**’** surface.

## CONCLUSION

The two DILP2:dmIR-ECD complexes described here underline the common site 1- binding motives for DILP2 and DILP5 hormones. One of the most unexpected results here is a low DILP2:dmIR-ECD occupancy: one (site 1) in conformer **1**, and two (site 1 and an in-between site 1**’**-2) in conformer **2**, despite very high 40:1 excess of the ligand over the receptor needed for the complexes, and with both conformers showing similar quaternary structures (**Fig. 3c**) and ligand binding modes. It seems that the N-/C-termini differences between DILP2 and DILP5 do not dictate unique allosteric effects through the dmIR. All this may indicate that the functional divergence of DILPs can be achieved – besides their distinct spatial and temporal expression patterns - not by significantly different structural receptor activation modes/signatures of DILP1-7, but by variations of their *k*_on_/*k*_off_ rates, and subsequent different duration of the signalling and endocytosis of the complexed dmIR, as already suggested (36) (39).

Whether the intercalation of DILP2 B22-B24 chain between L1β2 and α-CT**’** does really occur must be confirmer further, but such weakening of this crucial interface by the C-terminus of the B-chain presents an interesting alternative to the modulation of hormone:receptor binding and activation; especially in the biological environment where there is one IR-like receptor working with multiple insulin-like hormones (as in *Drosophila*).

In general, although DILP2:dmIR-ECD complex does not show very significant differences from DILP5:dmIR-ECD assembly, it revealed several original features, e.g., variability of the lower arm conformations, and possible alternative runs of the B20-B24 chains, that enrich our understanding of this system. It can also be envisaged that – serendipitously – DILP2 conformer **2**-DILP2 conformer **1**-DILP5 complex, together with a very dynamic character of conformer 2, represent some initial steps in the dmIR activation. Here, the arm-down DILP2 in conformer **2** can reflect the initial DILP2 binding (not committed to any site), on its way to the upper arm site 1, while conformer **1** is the active (‘stable’) form of the receptor (**Fig. 3c**). DILP5 complex would be then either a ‘supersaturated’ state of the dmIR, or a conformer characteristic for DILP5-driven receptor activation.

It is also interesting that the lack of symmetrical T-like DILP2 and DILP5 dmIR-ECD complexes suggest that this invertebrate Insulin Receptor-like system rely more on hIR - hIGF-1R allosteric ‘hybrid’ of signal transduction. It would mean, that a general architecture of human insulin:hIR binding is maintained in the dmIR system, but which still shows (in contrast to hIR) – at the supraphysiological level of the hormones – persistent negative cooperativity that prevent formation of a symmetrical T-like dmIR conformer. However, this still must be resolved by other DILPs:dmIR complexes.

## MATERIALS AND METHODS

### Production of dmIR-ECD

The *Drosophila melanogaster* insulin receptor ectodomain (dmIR-ECD) comprising of the native signal sequence of dmIR (264H-NYSYSPGISLLLFILLANTLAIQAV-290V) and DNA encoding the amino acids 264-1309 of dmIR - was expressed using the baculovirus-insect cell system with minor modifications from protocol as previously described by Viola et al. (40). Briefly, the codon optimized ectodomain (gift from Novo Nordisk) was sub-cloned into the PBac-4x-1 shuttle vector (Novagene) downstream of p10 promoter with the addition of start codon at the N-terminus and strep-tag-II at the C-terminus. The bacmid construct (gift from Ian M. Jones, University of Reading) was transformed into XL10-Gold ultracompetent cells and purified using the NucleoBond Xtra Midi EF kit (Macherey-Nagel). Prior to transfection, the bacmid was linearized with the BsuI restriction enzyme (NEB).

P1 viral stock was generated using adherent cultures in a 6-well format. Freshly split ExpiSf9 cells were seeded at 0.5×10^5^ cells/mL in 2 mL of fresh media and incubated at room temperature for 1 hour to allow cell attachment. The bacmid, shuttle vector, and transfection reagent (FuGENE® HD; Promega) were mixed at a ratio of 3:4:12, respectively, and diluted in 200 μL of fresh media. The transfection mixture was gently mixed and added dropwise to each well. Cultures were incubated at 28 °C for 5–7 days. The shuttle vector included eGFP as a reporter to monitor transfection efficiency.

P1 to P2 baculovirus amplification was performed in Sf9 suspension culture. ExpiSf9 cells were seeded at 1×10^6^ cells/mL and incubated at 28 °C with shaking (90 rpm, 50 mm orbital diameter). On the following day, when the cell density reached approximately 2×10^6^ cells/mL with >95% viability, 100 μL of P1 stock was added per 30 mL culture. Cultures were incubated at 28 °C for 4–5 days or until eGFP expression exceeded 80%.

### Expression and purification of dmIR-ECD

dmIR-ECD was expressed in 400 mL suspension cultures using 2 L Corning Erlenmeyer shaker flasks. One day prior to transfection, cultures were seeded at 1 × 10^6^ cells/mL and incubated at 28 °C with shaking. The following day, cultures were infected with baculovirus and monitored for eGFP marker expression. Expression typically exceeded 90% within 48 hours.

Conditioned media were harvested by centrifugation at 500 × g for 10 min, followed by 5,000 × g for 30 min. The supernatant was concentrated using tangential flow filtration (TFF) with a 30 kDa cut-off membrane (Repligen, S02-E030-05-N; 790 cm^2^ surface area), and buffer-exchanged into Buffer A (25 mM Tris-HCl, pH 7.4, 400 mM NaCl).

The diafiltrate was mixed with a precipitation solution containing NiCl_2_ and CaCl_2_, then centrifuged at 18,000 × g for 1 hour to remove precipitated debris. The clarified supernatant was loaded onto a 5 mL Strep-Tactin XT column (Cytiva), washed with 5 column volumes of Buffer A, and eluted with Buffer B (Buffer A supplemented with 50 mM biotin). Eluted fractions were pooled and concentrated using a 30 kDa cut-off Vivaspin concentrator (GE Healthcare), then further purified by size-exclusion chromatography (SEC) on a Superdex 200 Increase 10/300 column (Cytiva) in SEC buffer (20 mM HEPES, pH 7.4, 200 mM NaCl). Fractions containing dmIR-ECD were pooled and concentrated to 2 mg/mL using a 30 kDa Vivaspin concentrator.

### Synthesis of DILP2

DILP2 was prepared by total chemical synthesis using orthogonal protection of the sulfhydryl (–SH) groups of individual cysteine residues, following the protocol described by Liu et al. (41). The peptide chains were synthesized on a Spyder Mark IV Multiple Peptide Synthesizer (European Patent Application EP17206537.7), developed at the Development Center of the Institute of Organic Chemistry and Biochemistry of the CAS (DCIOCB, https://developmentcenter.group.uochb.cz/en). Fmoc-Asn(Trt)-Wang resin was used for chain A, and Fmoc-His(Trt)-Wang resin for chain B. Chain A comprises 26 amino acid residues and contains four cysteines, with each one protected with a distinct protecting group: CysA7–StBu, CysA8–Acm, CysA12–Mmt, and CysA21–Trt.

#### Chain A synthesis

Cysteines at positions A11 and A16 (Fmoc-Cys(StBu)-OH and Fmoc-Cys(Mmt)-OH, respectively) were manually attached to the resin. Each condensation reaction was performed using 5 equivalents of the protected amino acid in DMF, 0.45 M HOBt, 5 equivalents of HBTU (0.4 M in DMF), and 10 equivalents of DIPEA (1 M in DMF), with a reaction time of 2 hours. Coupling efficiency was monitored using the Kaiser test. Fmoc deprotection was carried out with 20% piperidine in DMF (5- and 20-minute treatments).

The remaining amino acids were coupled using automated synthesis. Each residue was introduced via a two-step coupling strategy. The first coupling employed 5 equivalents of the protected amino acid in DMF with 0.45 M HOBt and 5 equivalents of DIC (1 M in DMF), followed by a second coupling using HBTU, HOBt, and DIPEA under the same conditions as described above. Fmoc groups were deprotected as described above.

After completion of the synthesis, the resin was treated with 25% β-mercaptoethanol in DMF for 6 hours at room temperature. The treatment was then repeated overnight, followed by an additional 8-hour incubation with 20% tributylphosphine in DMF containing 2% H_2_O (v/v) to ensure complete deprotection of the StBu group. To confirm the removal of the StBu protecting group from the cysteine at position A11, a small sample of the resin was cleaved with TFA and analyzed by LC-MS.

Subsequently, the resin was washed with DMF and DCM, then treated with 10 equivalents of 2,2′-dithiobis(5-nitropyridine) (DTNP) in DCM for 1 hour at room temperature. Following the reaction, the resin was washed again with DMF and DCM, and subsequently treated with 1% TFA and 5% TIS in DCM (5 × 2-minute treatments at room temperature). Finally, the resin was washed once more with DMF and DCM and stirred in DCM for an additional hour at room temperature.

Final cleavage of the peptide from the resin was performed using a TFA/TIS/H_2_O mixture (95:2.5:2.5, v/v/v) for 1.5 hours. The crude peptide was precipitated by the addition of cold diethyl ether and subsequently purified by reversed-phase HPLC on a Nucleosil 100-7 C8 column (250 × 10 mm, 7 μm, Macherey-Nagel) using a Waters HPLC system (Waters 600 with 2487 Dual λ Absorbance Detector) at a flow rate of 4 mL/min. Separation was achieved using a gradient of acetonitrile in water with 0.1% TFA: solvent A = 0.1% TFA (v/v) in H_2_O; solvent B = 0.1% TFA (v/v) in 80% CH_3_CN; gradient: t = 0 min, 10% B → t = 30 min, 100% B.

The purity of the final peptide was assessed by analytical HPLC using a Waters™ e2695 system with a 2489 UV/Vis Detector and a YMC-Triart Bio C4 column (250 × 4.6 mm) at a flow rate of 1 mL/min, employing the same gradient and solvent system as described above. Compounds were detected at 218 nm.

#### Chain B synthesis

The B-chain (CysB3–Acm, CysB15–Trt) was synthesized using standard Fmoc-based solid-phase peptide synthesis (SPPS), following the procedure described above. Cleavage from the resin was performed using a cocktail of TFA:TIS:H_2_O (95:2.5:2.5, v/v/v) in the presence of 15 equivalents of DTNP for 2 hours at room temperature. The crude peptide was then isolated, purified, and characterized using the same protocols as described for the A-chain.

#### Ligation of chains

The A- and B-chains of DILP2 were combined in 6 M urea and 0.2 M NH_4_HCO_3_ (approximately 1:1 molar ratio) buffer (pH 8.0). The mixture was stirred at room temperature until fully dissolved. After 5 minutes, 25 equivalents (relative to chain A) of freshly prepared iodine solution in AcOH were added. The reaction mixture was stirred gently for an additional 10 minutes at room temperature. To quench the reaction, a 1 M solution of ascorbic acid was added dropwise until the solution became colorless. The resulting solution was diluted with H_2_O, and the peptide was purified by reversed-phase HPLC as previously described. The purity of the final product was confirmed by RP-HPLC (**SI Figure 1a**), and the correct molecular weight of DILP2 was verified by high-resolution mass spectrometry (LTQ Orbitrap XL, Thermo Fisher Scientific, Waltham, MA) (**SI Figure 1b**).

### Grid Preparation and data collection

DILP2 was freshly dissolved in 0.1 percent acetic acid to a final concentration of 10mg/ml which was subsequently diluted in SEC buffer (25mM HEPES pH 8.0, 200mM NaCl) to a final concentration of 1mg/ml working stock. The dmIR-ECD was diluted to 0.2mg/ml in SEC buffer and incubated with 40-fold molar excess of DILP2 at room temperature for 30 mins. 3 μL of the mixture were applied to UltrAuFoil R 1.2/1.3 300 mesh grids glow-discharged using a PELCO easiGlow glow-discharge at 20 mA, 0.38 mBar and vitrified using a using the Mark IV Vitrobot (Thermo Fisher Scientific) at 4 °C and 95% relative humidity for 2 s with −10 blot force. Screening and data collection were performed on Thermo Fisher Scientific Glacios Cryo-TEM with Falcon 4 detector. Automated data collection was performed using Thermo Fisher EPU software. Two datasets were collected from different grids with 17397 micrographs in total (4397 and 13000). For both datasets, movies were collected with pixel size 0.574Å/pixel with target defocus range of 0.2-2 μm. Each movie was recorded with a total fluence of 50.0 e^−^/Å^2^ over 2.77 s, corresponding to a flux of 6 e^−^/pixel/s (**SI Table 1**).

### Data processing and Model reconstruction

Both datasets were processed using Relion 4.1 (42) and analysed in ChimeraX (43). For dataset 1, micrographs were initially manually sorted to eliminate empty or icy ones; resulting in 1404 micrographs out of the initial 4397 (see **SI Figure 2** for the project flowchart). Movies recorded in EER format were motion corrected using RELION’s implementation of the MotionCor2 algorithm (42, 44). CTF parameters were estimated using CTFFIND4.1 (45). Subsequently micrographs recorded at >2 μm defocus or with estimated astigmatism >600 Å and CTF fit resolution > 10Å were discarded to further remove bad images, with 1068 micrographs remaining. Particle picking was performed with Topaz (46) from a subset of 100 micrographs with the Topaz default model finding 38812 particles. Particles with autopicking figure of merit (FOM) ≥ 0 were extracted in 480 pixel boxes and down-sampled to 2.296 Å/pixel, resulting in 12065 particles. After 2D classification the best classes (5798 particles) were selected to train Topaz model. The trained model was used to pick 72748 particles from the complete dataset. Further 2D classification was carried out to select classes to create the initial model (17 classes were selected with not more than 500 particles per class, with final 8267 particles). This initial model was used as a reference in 3D classification, that was carried out for 44323 particles selected in a separate 2D classification job. The best class (13103 particles) was selected for 3D refinement, the resulting map was used as a reference in the next round. In round 2, 13000 movies from the second dataset were added; these were motion corrected and CTF parameters estimated as described for round 1. This time there was no manual selection, but micrographs collected at >2 μm defocus or with estimated astigmatism >600 Å were discarded, resulting in 12103 micrographs (**SI Figure 2**). The best 3D class from round 1 was used for training Topaz model, followed by particle picking. 616516 particles with auto-picking FOM ≥ -1 were extracted from those micrographs in 480 pixel boxes and down-sampled to 2.296 Å/pixel. Two rounds of 2D classification were carried out to clean the dataset, resulting in 146 595 particles. 3D classification was carried out using the final model from the 1st dataset initially low pass filtered to 20 Å as the reference, followed by 3D refinement, achieving 5.7Å resolution. This map was used as a reference in the third round. In round 3, more particles, with FOM ≥ -3, were extracted from round 2 topaz picking resulting in 882882 particles. This was followed by 25 iterations of 3D classification, and further 25 iterations with 0.9 degrees sampling and local searches (+/-5.625 degrees). Further 3D refinement with Blush regularization (47) was carried out for 386187 particles from the best class resulting in 4.66Å resolution reconstruction. This was followed by Bayesian polishing with shiny particles extracted from movie frames in 648 pixel boxes and down-sampled to 1.29 Å/pixel (final box size 288 pixels), CTF refinement (48), and another 3D refinement resulting in 3.95Å resolution. After a further 3D classification with local searches (**SI Figure 2**) the best-looking class was chosen, with 127081 particles. Final 3D refinement resulted in overall 3.75Å resolution, although the worst defined regions have resolution below that value (**SI Figure 3**).

An alternative route of particle analysis lead to the model two – conformer **2**, with two DILP2 bound states. For this, the particles were re-extracted with FOM of 0.05. The extracted particles were cleaned in several iterations of 2D classifications and subset selections. The resulting final subset of 131,611 particles was used in generating the initial model and followed by 3D classification to explore heterogeneity. The generated 3D classes were analyzed in ChimeraX for structural heterogeneity. The particles for model with additional density (57300 particles) were extracted and were further used in high resolution refinements, as described above.

### Model building and refinement

For both structures, cryo-EM structure of dmIR-ECD bound with DILP5 (PDB ID 8CLS) (40) was used as the initial model. For the model with one DILP2 per monomer – conformer **1**, the model was fitted into the 3.75 Å resolution map using the rigid body fitting algorithm “fit in map” implemented in ChimeraX (43). Further model building was carried out in Coot (49). The models of individual domains were calculated using AlphaFold2 (50), and these domains were first superposed on the corresponding domains of the 8CLS model, and then fitted with jiggle fit option. DILP2 molecule was generated using homology modelling implemented in Modeller (51), and structurally aligned with DILP5 molecules using matchmaker function in ChimeraX (and later verified and corrected where necessary using also the AlphaFold2 model of DILP2 as well). Real-space refinement was carried out in ISOLDE (52) alternated with reciprocal space refinement with REFMAC5 (CCP4 suite, version 7.1, ccpEM version 1.5) (53). Visible N-glycans were added in Coot (49).

For the model with two DILP2 molecules – conformer **2**, the second DILP2 was manually placed and fit to the 6.25 Å resolution map in ChimeraX using the fit to map function. The fitted model was used in molecular dynamics flexible fitting method to obtain optimal refinement using the online server Namdinator (54). The model was refined using Isolde (52) and was further refined in Phenix for side chain glycans (55). Both models were validated using Compehensive cryoEM validation tool in Phenix including MolProbity (56) (**SI Table 1**). The LSQ and SSM superpositions have been done in Coot (49). Figures have been made in ChimeraX (43) and CCP4MG programme (57).

## DATA AVAILABILITY

The atomic coordinates of the DILP2:dmIR-ECD conformer **1** and conformer **2** (and corresponding cryo-EM density maps) have been deposited in the Protein Data Bank and EMDB data banks with the accession codes: 9HKT (DOI: https://doi.org/10.2210/pdb9HKT/pdb), and 9HNI (DOI: https://doi.org/10.2210/pdb9HNI/pdb), respectively, and EMD-52236 and EMD-52310.

## ETHICS DECLARATION

The authors declare no competing interest, and this work did not require ethical approval from a human subject or animal welfare committee.

## CONTRIBUTIONS

**TS** produced dmIR-ECD, and worked on data acquisition, cryo-EM structure solution, model building/refinement, **OM** worked on data acquisition, cryo-EM structure solution, model building/refinement, **HTJ, MC** and **MNI** helped with some cryo-EM aspects of this work, **ML** and **JJ** designed and performed synthesis of DILP2, **PDM** co-inspired the project, **AMB** designed the study, analysed the results and wrote the manuscript.

## ACKNOWLEDGEMENTS

**TS** and **AMB** were supported by BBSRC Grant BB/W003783/1, **OM** was funded by the G. G. Dodson Fund in York sponsored by Novo Nordisk A/S (Maaloev, Denmark), ML and JJ were supported by the project National Institute for Research of Metabolic and Cardiovascular Diseases (Program EXCELES, ID Project No. LX22NPO5104, Funded by the European Union-Next Generation EU), and by the Academy of Sciences of the Czech Republic (Research Project RVO:52, support to the Institute of Organic Chemistry and Biochemistry).

Novo Nordisk is greatly acknowledged for funding of the G. G. Dodson Fund in York. We thank the YSBL Cryo-EM Facility at the University of York (Glacios electron microscope funded by the Wellcome Trust (grant number 206161/Z/17/Z)) for access, and Dr Johan Turkenburg and Mr Sam Hart for assistance.

