## Supplemental Information for "Structural basis of *Drosophila* insulin receptor activation by DILP2 hormone"

### for

[±]deceased

**SI Figure 1.** HPLC analysis (a) and deconvoluted HR-MS spectrum (b) of synthesized DILP2. Chemical Formula: $C_{242}H_{397}N_{69}O_{77}S_8$, Exact Mass: 5757.70.

(***a***)

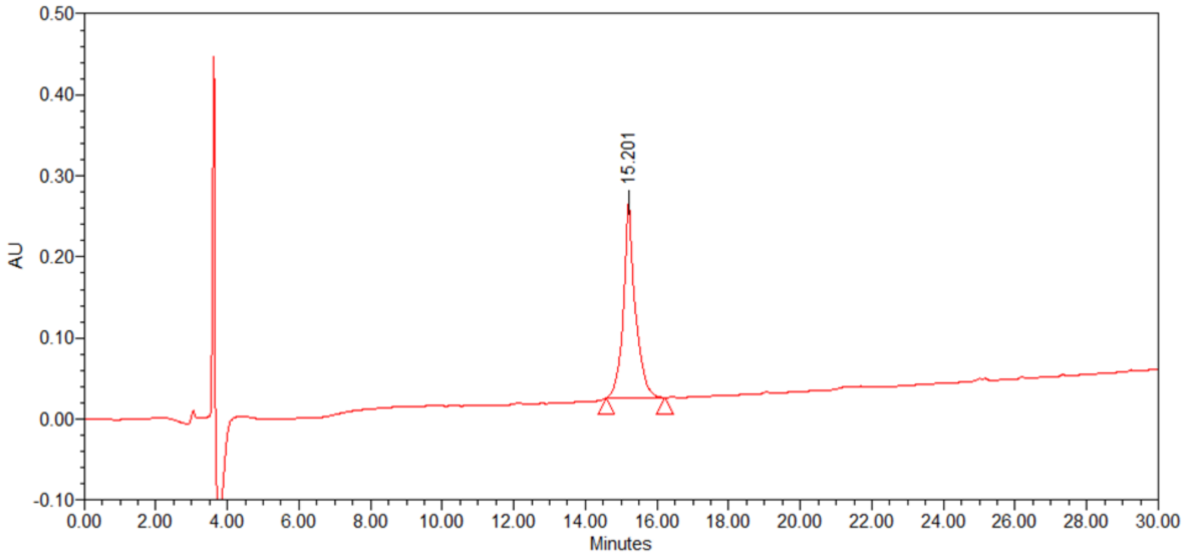


(***b***)

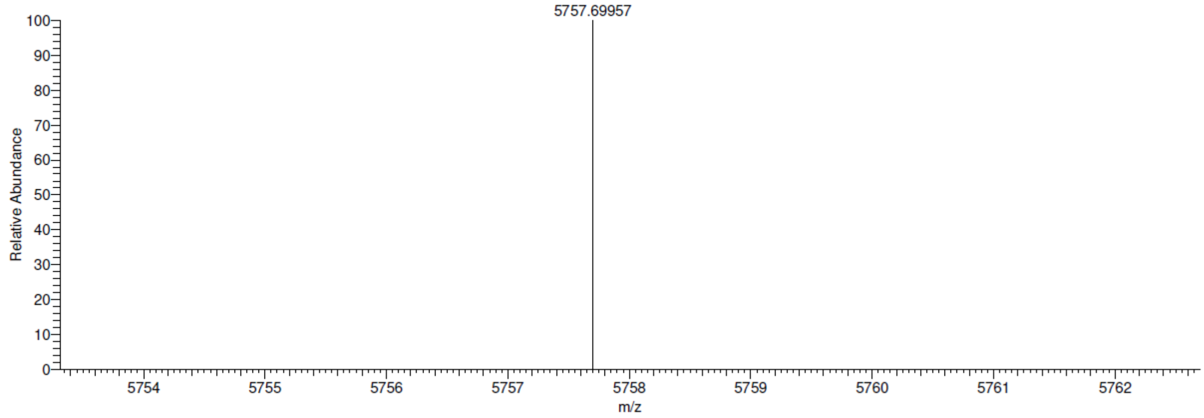

**SI Table 1.** Cryo-EM data collection parameters, refinement and validation statistics.

| **Data Collection** | | |
|---|---|---|
| Microscope/Detector | TFS Glacios / Falcon IV | |
| Voltage (kV) | 200 | |
| Magnification | 240K | |
| Pixel size (Å) | 0.574 | |
| Defocus range (µm) | 0.6-2.0 | |
| Total dose ($e^-/Å^2$) | 50 | |
| Flux ($e^-/Å^2/s$) | 6.27 | |
| Number of micrographs | 17397 | |
| Image Processing | | |
| Software | Relion 5 | |
| Total particles picked | 882882 | 257272 |
| Final particles used | 127081 | 57317 |
| Symmetry imposed | C1 | |
| Final map resolution (Å, FSC 0.143) | 3.76 | 6.25 |
| Sharpening B-factor ($Å^2$) | -66.0 | -144.784 |
| Model Building and Refinement | | |
| Initial model used | 8CLS, AF2 | |
| Refinement software | ccpEM, Refmac | |
| Model resolution (Å, FSC 0.5) | 4.2 | 7.8 |
| Non-hydrogen atoms | 14220 | 14354 |
| Average B-factor | 119.6 | 564.3 |
| RMSD bond lengths (Å) | 0.007 | 0.012 |
| RMSD bond angles (°) | 1.364 | 2.067 |
| Model Validation | | |
| MolProbity score | 1.91 | 2.55 |
| Clashscore | 3.45 | 7.99 |
| Ramachandran favoured (%) | 92.37 | 84.82 |
| Ramachandran allowed (%) | 7.63 | 15.18 |
| Ramachandran outliers (%) | 0 | 0 |
| Rotamer outliers (%) | 2.46 | 3.97 |
| Database Accession Numbers | | |
| EMDB ID | EMD-52236 | EMD-52310 |
| PDB ID | 9HKT | 9HNI |

**SI Figure 2.** Workflow and data processing.

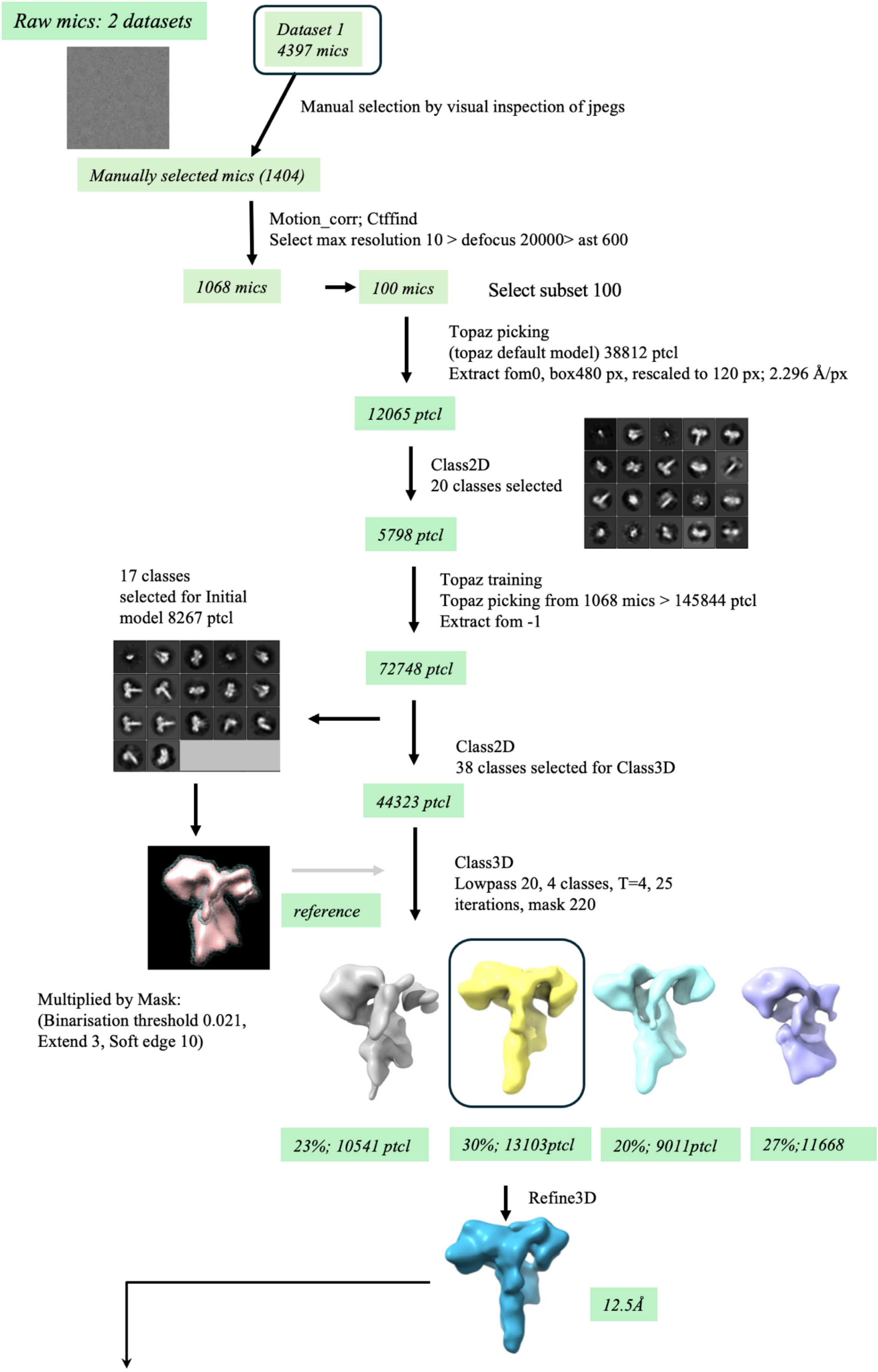

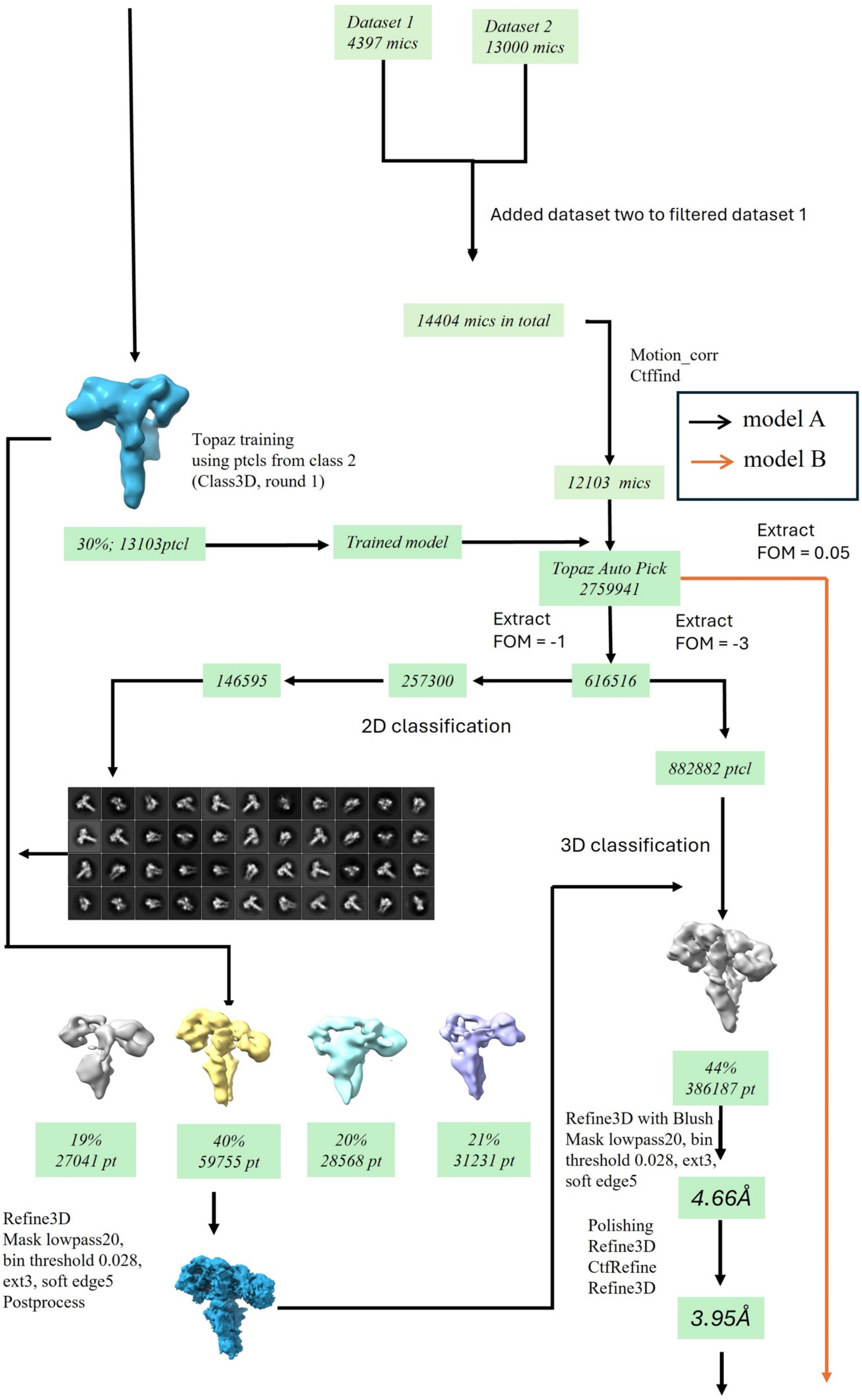
Dataset 1
4397 mics
Dataset 2
13000 mics
Added dataset two to filtered dataset 1
14404 mics in total
Motion_corr
Ctffind
model A
model B
12103 mics
Topaz training
using ptcls from class 2
(Class3D, round 1)
30%; 13103ptcl
Trained model
Extract
FOM = 0.05
Topaz Auto Pick
2759941
Extract
FOM = -1
Extract
FOM = -3
146595
257300
616516
2D classification
882882 ptcl
3D classification
44%
386187 pt
Refine3D with Blush
Mask lowpass20, bin
threshold 0.028, ext3,
soft edge5
4.66Å
Polishing
Refine3D
CtfRefine
Refine3D
3.95Å
19%
27041 pt
40%
59755 pt
20%
28568 pt
21%
31231 pt
Refine3D
Mask lowpass20,
bin threshold 0.028,
ext3, soft edge5
Postprocess

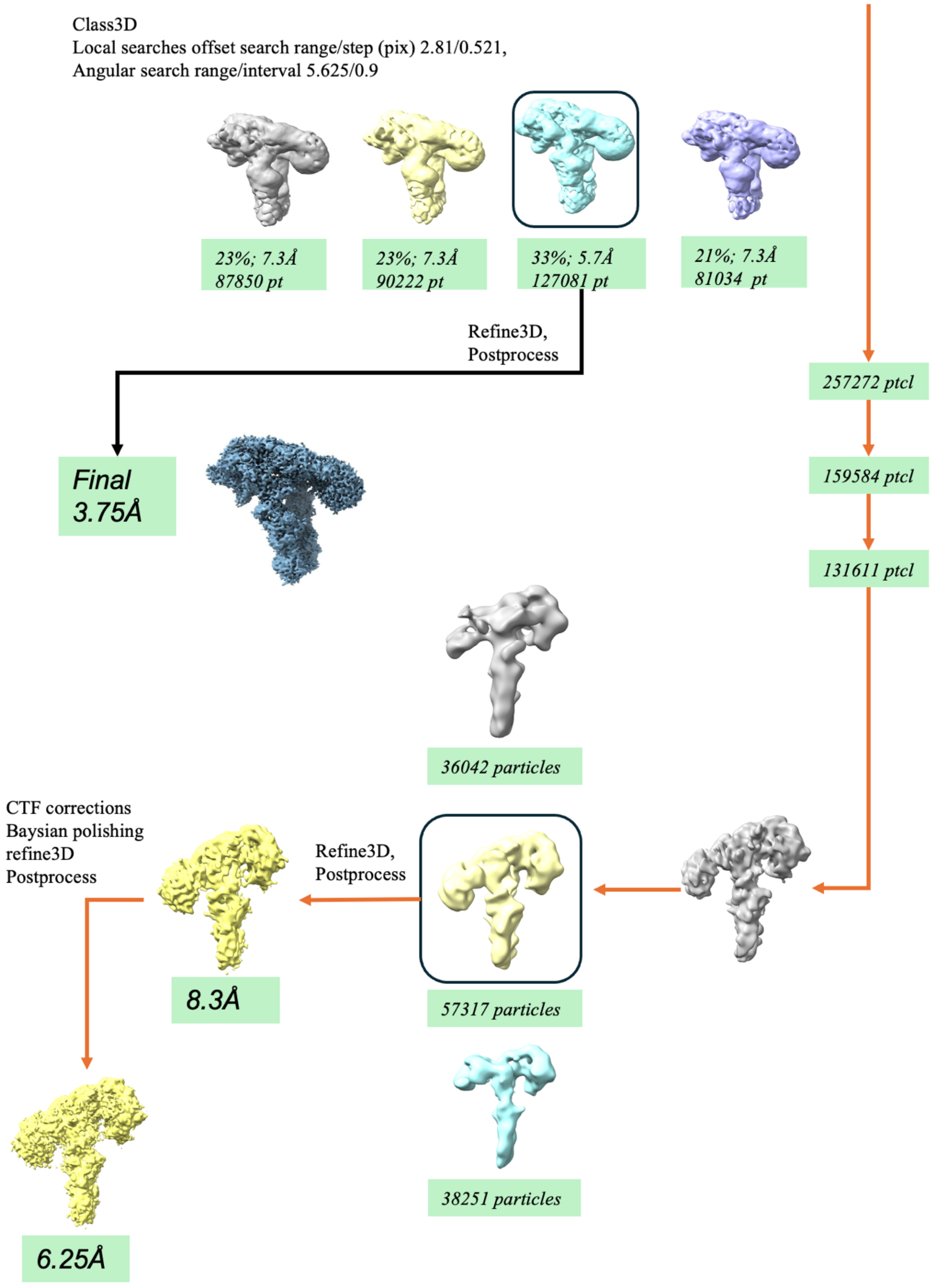
Class3D
Local searches offset search range/step (pix) 2.81/0.521,
Angular search range/interval 5.625/0.9
23%; 7.3Å
87850 pt
23%; 7.3Å
90222 pt
33%; 5.7Å
127081 pt
21%; 7.3Å
81034 pt
Refine3D,
Postprocess
257272 ptcl
159584 ptcl
131611 ptcl
Final
3.75Å
36042 particles
CTF corrections
Baysian polishing
refine3D
Postprocess
Refine3D,
Postprocess
8.3Å
57317 particles
38251 particles
6.25Å

**SI Figure 3.** (a) Curves of FSC computed for phase-randomized masked half-maps (red), unmasked half-maps (green), masked half-maps (blue) and the corrected FSC curve (black). (b) CryoEM-density map filtered and coloured according to local resolution.

a

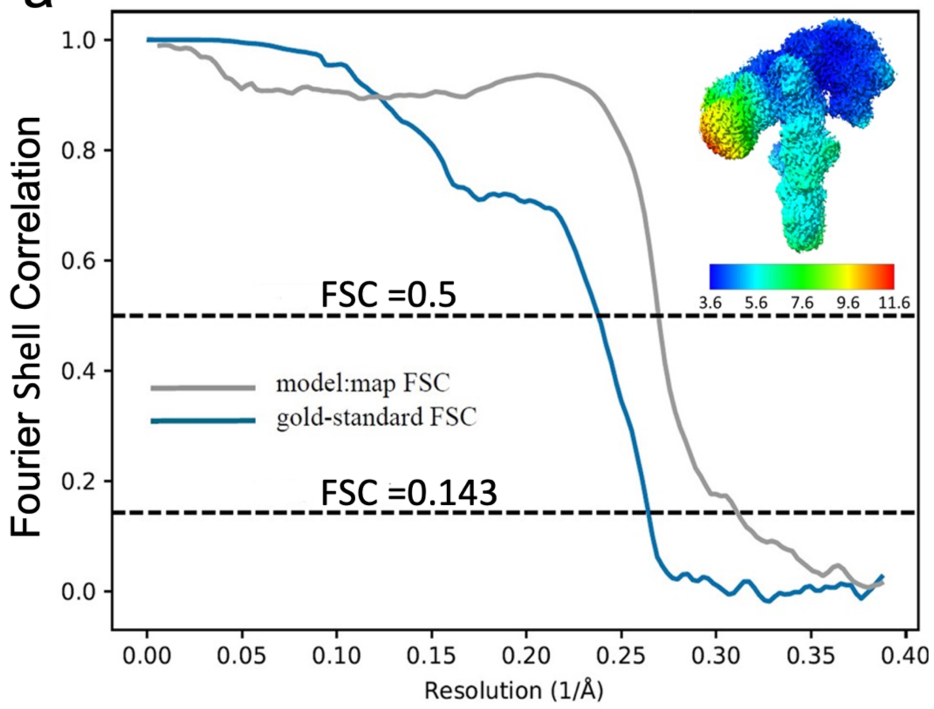


b

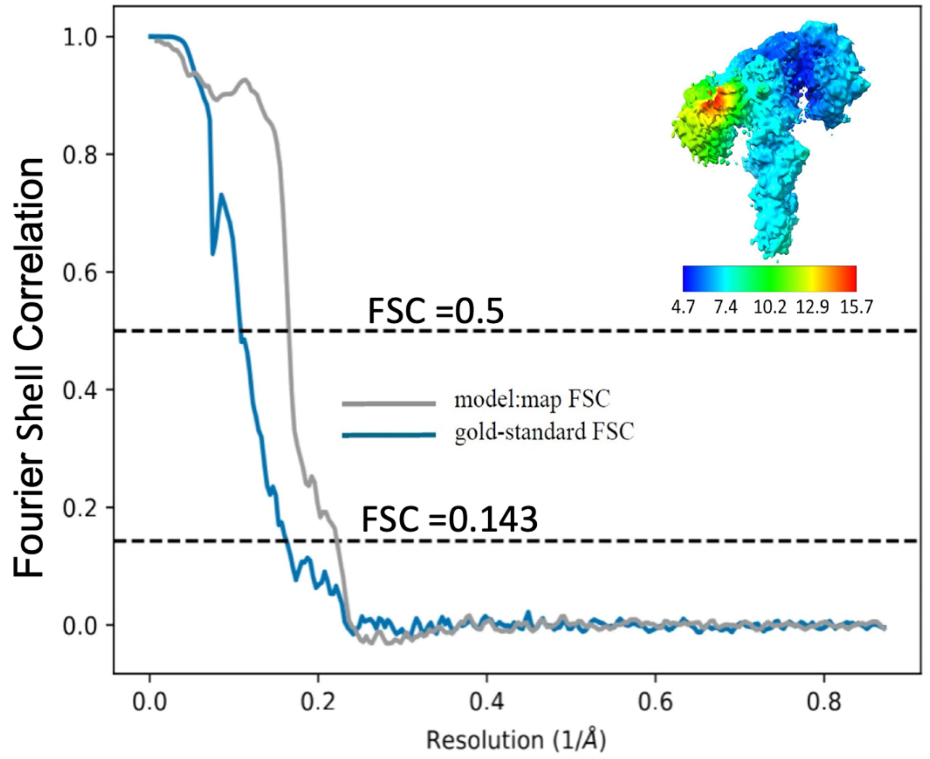

**SI Figure 4.** Cryo-EM density maps and models for site 1 in conformers **1** (**a**) and **2** (**b**) , and second DILP2 in between sites 1'-2 in conformer **2** (**c**).

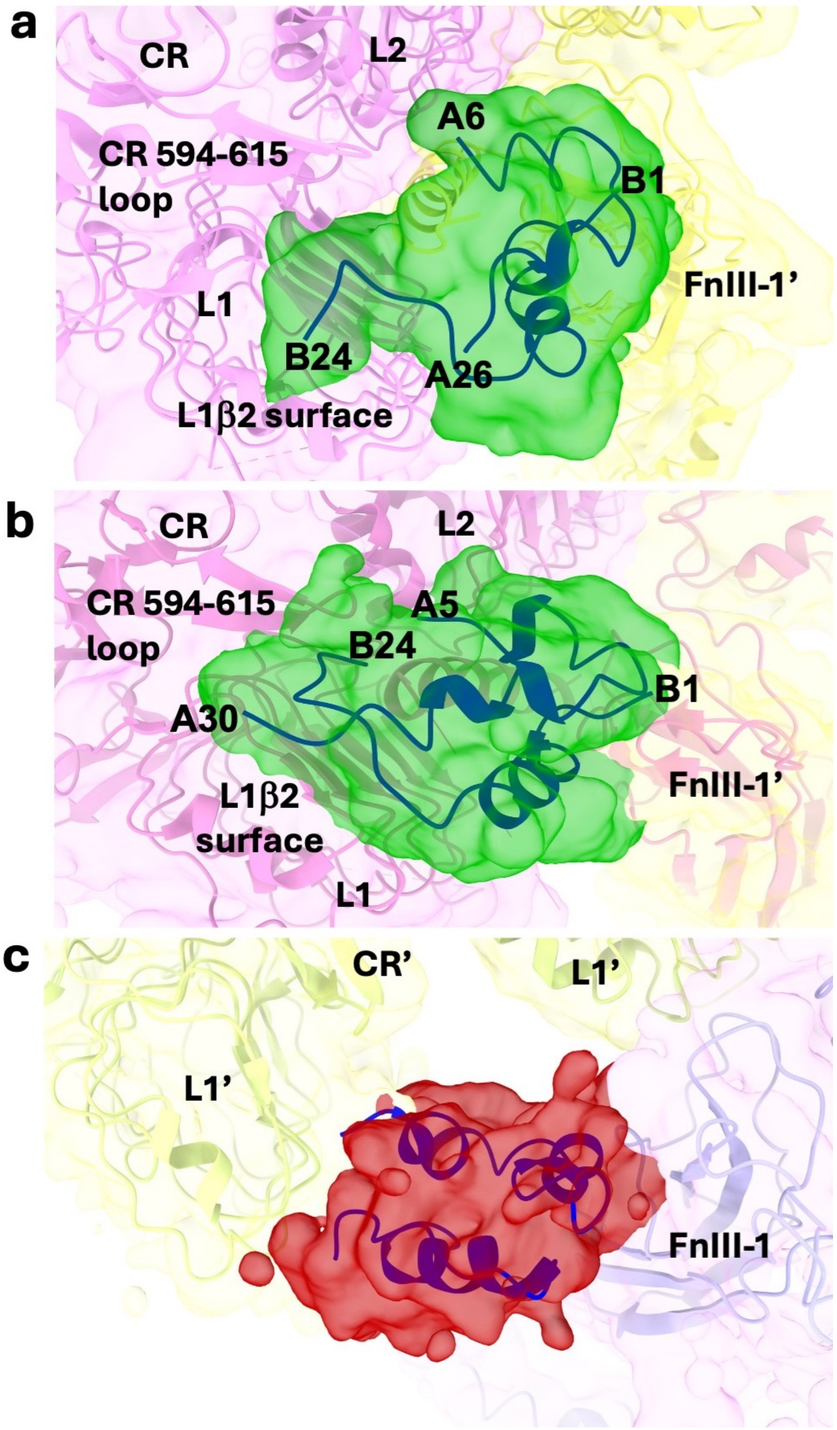
